# Effects of tachykinin-related peptides on the reproductive system of *Tenebrio molitor* females: implications for insect breeding and pest control

**DOI:** 10.1101/2025.08.25.672133

**Authors:** N. Konopińska, K. Walkowiak-Nowicka, Sz. Chowański, G Nowicki, A. Urbański

**Affiliations:** Department of Animal Physiology and Developmental Biology, Faculty of Biology, Adam Mickiewicz University, Poznan, Poland; GenXone S.A., Złotniki, Poland

**Keywords:** insect reproduction, vitellogenin, neuropeptides, insect mass rearing, insecticides

## Abstract

The global population, which is expected to reach 10.4 billion by 2086, will significantly increase the demand for sustainable food sources. Edible insects such as *Tenebrio molitor* are promising alternatives because of their nutritional value, low environmental footprint, and suitability for mass rearing. However, the efficiency of industrial production depends on the optimization of reproductive processes. Moreover, *T. molitor* is also a pest species that contributes to grain loss, highlighting the dual need for strategies that increase reproduction under farming conditions and suppress fertility in pest populations.

Neuropeptides, including tachykinin-related peptides (TRPs), are known regulators of metabolism and immunity, but their role in reproduction remain unclear. Here, we investigated whether TRPs are involved in female *T. molitor* reproduction. Expression analyses revealed strong correlations between *TRP*, *TRPR* and *vitellogenin* (*Vg*) gene expression, suggesting TRP-mediated stimulation of yolk precursor synthesis. The application of Tenmo-TRP-7 affects basic reproductive parameters, including egg production, follicular epithelium permeability, and terminal oocyte volume. These effects are confirmed by the use of dsRNA directed against the gene encoding TRP precursor.

These findings show that TRPs regulate reproduction at multiple levels, positioning them as molecular targets for both enhancing insect farming and developing environmentally safe pest control strategies.

## 1. Introduction

Despite assurances from many scientific centers about the coming stabilization of the population, the global population continues to grow. According to United Nations projections, the population peak in 2086 will reach approximately 10.4 billion people ^1^. The growing population is associated with an ever-increasing demand for food - according to a meta-analysis published in Nature Food, this demand could increase by up to 56% by 2050 ^2^. Meanwhile, traditional animal husbandry is putting a significant strain on the environment, resulting in high water consumption, greenhouse gas emissions and ecosystem degradation ^3^. Faced with these challenges, the search for sustainable, efficient, and safe sources of protein and other nutrients has become crucial. One of the most promising solutions are edible insects, such as the mealworm beetle (*Tenebrio molitor*), which was approved by the European Union for consumption. This species has a high nutritional value - it contains high amounts of easily digestible protein and fats, including unsaturated fatty acids, and its amino acid profile meets FAO/WHO standards ^4^. In addition, larvae grow quickly, are easy to raise and require far fewer resources than livestock - they use less water, feed and space, while emitting fewer greenhouse gases ^5–7^. They can also be fed on organic waste, which fits in with the tenets of a closed-loop economy. Its safety as a food ingredient has been confirmed by the European Food Safety Authority (EFSA) ^4^. However, in order to fully realize the potential of *T. molitor* in food production, it is necessary to optimize its reproductive cycle, which determines the efficiency of breeding on an industrial scale.

On the other hand, *Tenebrio* is also an insect that causes serious economic losses. Those insects are responsible for the destruction of up to 15% of global losses in grain products ^8,9^. Due to the huge range of losses generated by these insects, pesticides are most often used to control number of insects, but despite their effectiveness, commonly used chemical insecticides have numerous drawbacks - they can harm pollinators, humans and other non-target organisms and contribute to the loss of biodiversity ^10^. In addition, insects develop resistance to insecticides over time, resulting in the need to increase insecticides doses. In 2022, insecticide consumption reached as high as 3.75 million tons, according to the FAO ^11^. In the face of these challenges, there is a growing search for new, selective methods of crop protection that are safe for the environment and non-target organisms.

In both cases interest in insect neuropeptides and their signalling pathways as a new class of compounds that can support both breeding and plant biodefense is increasing ^12^. Particularly promising are tachykinin-related peptides (TRPs) which regulate many processes related to behavior, development and growth, including metabolism, immunity and food intake ^13–15^. However, their effects on the reproductive system of insects, especially *T. molitor*, remain poorly understood. Here we show that TRPs can influence the activity of the reproductive system. We demonstrated a significant correlation between the expression of genes encoding TRPs and TRPRs and the expression of gene encoding vitellogenin (Vg), a yolk precursor protein. We show that TRPs also influence key reproductive parameters, including egg number and changes in ovarian structure. Their direct effect was confirmed via RNA interference (RNAi) method with dsRNA knockdown of *TRP*, gene encoding TRP precursor.

By identifying the reproductive functions of TRPs and confirming their direct effects via RNAi, we provide a foundation for the targeted modulation of reproductive processes in *T. molitor*, which may significantly increase the efficiency and scalability of industrial insect farming. At the same time, the ability to interfere with TRP signalling pathways opens new possibilities for controlling the fertility of pest species in a selective and environmentally responsible manner. This dual potential highlights the broader applicability of insect neuropeptides as molecular tools for both sustainable food production and next-generation plant protection strategies.

## 2. Results

### 2.1. Expression levels of *TRP*, *TRPR* and *Vg* in 1-7-day old females of *Tenebrio molitor*

Females of *T. molitor* reach sexual maturity and lay their first eggs as early as the fourth day after eclosion, making the period from day 1 to day 7 of adulthood particularly important for the regulation of reproductive processes ^16^. Following eclosion, females undergo dynamic physiological changes, including ovarian maturation and metabolic remodelling, which are mediated by numerous hormones ^16^. One of the key elements of these processes is vitellogenin, a glycoprotein synthesized in the fat body and transported via the hemolymph to the ovaries, where it serves as a yolk precursor stored in the oocytes ^17^. To determine whether TRPs may be involved in regulating reproductive system activity in females, it was first necessary to analyse the expression levels of genes encoding both the TRP precursor and its receptor, as well as Vg. This enabled the evaluation of potential relationships between neuropeptidergic regulatory mechanisms and ovarian maturation and yolk production during and after the first ovarian cycle. Due to correlation of obtained data with systemic changes of the TRP system, we also collected female heads. Since the insect head contains the brain and associated neural tissues, it serves as a representative sample for studying the nervous system ^18^.

The results show that *TRP* and *TRPR* expression in the head (*TRP* H and *TRPR* H) remains generally stable, with significant increases observed on day 5 (*TRP* H) and on days 6 and 7 (*TRPR* H). In the fat body, precursor gene expression significantly increases from day 3 to day 7, whereas *TRPR* expression increases from day 2 through day 7. Additionally, on days 6 and 7, a significant downregulation of *TRPR* gene was noted in the ovary. For *Vg* in the fat body (*Vg* FB), overexpression was observed on days 3–5 and day 7, and in the ovaries of 2- and 4-day-old females (Fig. 1A and Fig. S1).

**Fig. 1.**
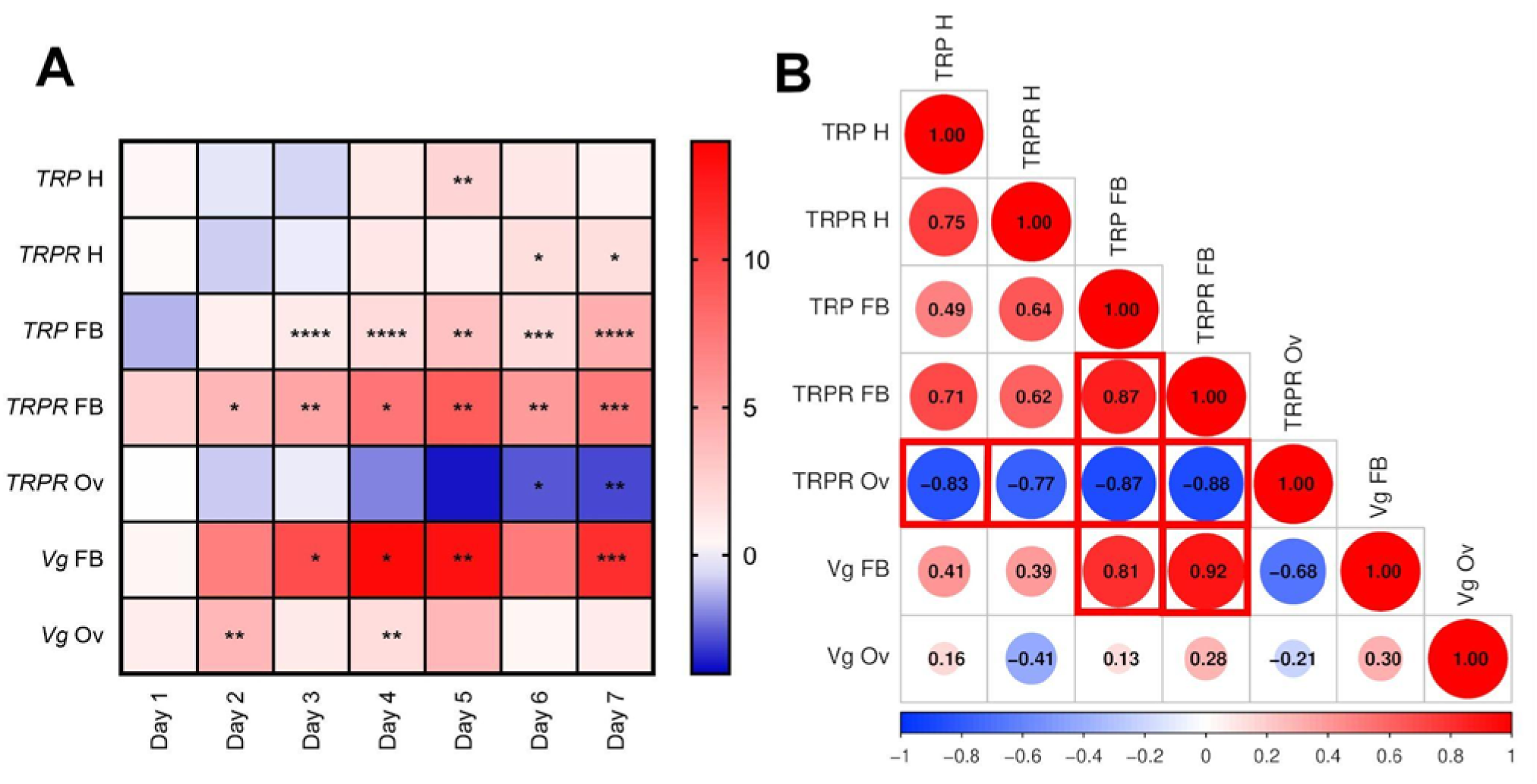
A - Heatmaps showing the changes in the expression levels of genes encoding TRP precursor, TRP receptor (TRPR) and vitellogenin (Vg) in the head (H), fat body (FB) and ovaries (Ov) in 1-7-day old females of *Tenebrio molitor*. The values are expressed as log2fold values, shades of red indicate upregulation, shades of blue indicate downregulation, \**p* ≤ 0.05, \*\**p* ≤ 0.01, *** *p* ≤ 0.001, **** *p* ≤ 0.0001 (compared to samples collected on Day 1). B - The correlation matrix showing the dependencies between the expression levels of *TRP* and *TRPR* and *vitellogenin* (Vg) genes in the head (H), fat body (FB), and ovaries (Ov). To estimate the correlation of the data, the Pearson correlation coefficient method was used. The matrix was generated via SRplot software (https://www.bioinformatics.com.cn/srplot). Size of the dot - level of the *r* value. The *r* value is presented in the middle of the dot. Different colors indicate different *r* values. Red shading indicates positive correlations (r ≥ 0), and blue shading indicates negative correlations (r ≤ 0). Red squares indicate statistically significant correlations (*p* ≤ 0.05).

Additionally, a correlation analysis was performed to determine whether there were dependencies between the expression of the studied genes (Fig. 1B). The results indicate a significant positive correlation between *TRP* and *TRPR* in the fat body, as well as between *Vg* and *TRP* and *TRPR* in this tissue. Interestingly, a significant negative correlation was observed between the expression of *TRPR* in the ovaries and the expression of *TRP* and *TRPR* in the head and fat body.

## 3. Evaluation of TRP on female reproduction

Given prior findings demonstrating dynamic changes in *TRP* gene expression during the adult stage, we next aimed to investigate its functional role in female reproductive physiology. Therefore, we evaluated the effects of both exogenous Tenmo-TRP-7 (at concentrations 10^-7^ M and 10^-5^ M) application and RNAi knockdown of *TRP* gene (ds*TRP*) on adult females. The neuropeptide concentrations were selected based on previous studies investigating insect neuropeptides, where similar ranges were shown to effectively elicit physiological responses without inducing toxic effects ^13^.

### 3.1.1. Survival experiment

In the survival experiment, Tenmo-TRP-7 was administered to females on days 1 and 4 of adulthood to examine the neuropeptide’s impact at different stages of the first ovarian cycle. Additionally, ds*TRP* was injected at days 1 and 4 post-eclosion, with survival monitored from day 8, when systemic RNAi effects are expected to be established (Fig. 7). The results show that application of Tenmo-TRP-7 does not affect female mortality at any of the tested time points. Similarly, no statistically significant changes in survival were observed following ds*TRP* treatment, indicating that manipulation of TRP signaling does not influence adult female survival under the tested conditions (Fig. 2S).

**Fig. 2.**
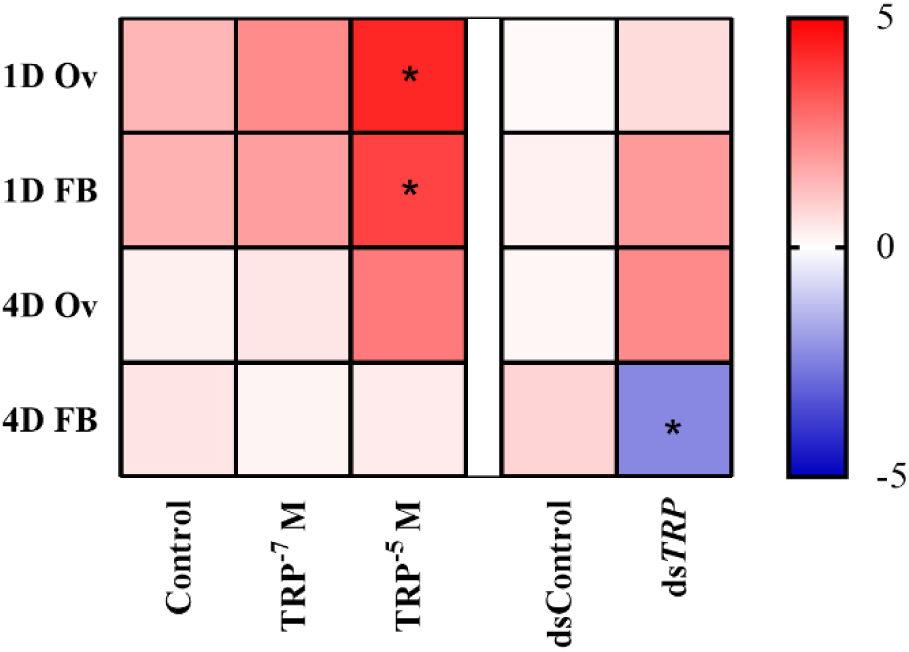
Heatmaps showing the changes in the expression levels of genes encoding vitellogenin in the fat body (FB) and ovaries (Ov) of 1-day-old (1D) and 4-day-old (4D) *T. molitor* females 24 h after the application of physiological solution (control) and Tenmo-TRP-7 at a concentration 10^-7^ M and 10^-5^ M. Additionally the *Vg* level was also tested after ds*GmLys* (dsControl) and ds*TRP* application. The values are expressed as log2fold values, shades of red indicate upregulation, shades of blue indicate downregulation, \**p* ≤ 0.05 (compared to Control groups).

### 3.1.2. Expression levels of gene encoding vitellogenin after the application of Tenmo- TRP-7 and dsRNA

The next step was to check whether Tenmo-TRP-7 or ds*TRP* application affects the *Vg* level in two key reproductive tissues – the fat body and ovaries.

The results revealed that significant upregulation of vitellogenin gene expression occurred only in 1-day-old females following Tenmo-TRP-7 injection at a concentration of 10^-5^ M, in both the fat body and ovaries. Interestingly, in 4-day-old females, silencing of the *TRP* gene resulted in a significant downregulation of *Vg* expression, but only in the fat body, with no observable changes in the ovaries (Fig. 2).

### 3.1.3. Changes in egg production

The observed correlations between the expression levels of *TRP*, *TRPR*, and *Vg* suggest a potential role of TRP in the regulation of the reproductive system. Since Tenmo-TRP-7 influences *Vg* expression, we investigated whether this effect also translates into changes in egg production. Initially, the effect of Tenmo-TRP-7 on egg production was examined in 1- and 4-day-old females. The results revealed a significant increase in the number of eggs laid by 1-day-old females following treatment with TRP at a concentration of 10^-5^ M (Fig 3A). In 4-day-old females, an increased number of eggs was observed after treatment with Tenmo-TRP-7 at a concentration of 10^-7^ M (Fig 3B). Silencing of the gene encoding the *TRP* did not lead to statistically significant changes in egg production in either age group (Fig. 3C).

**Fig. 3.**
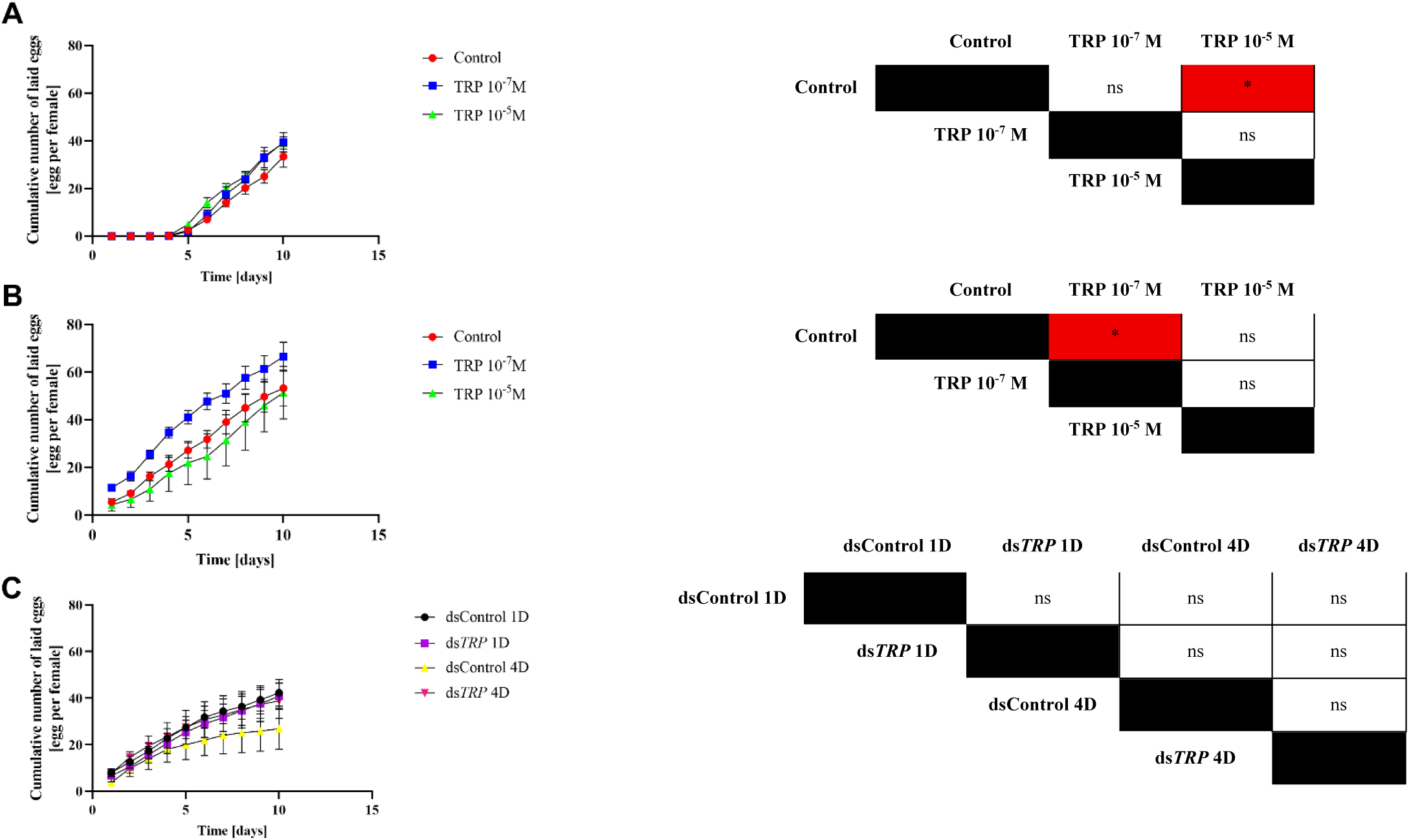
Cumulative number of eggs laid by 1-day-old (A), 4-day-old (B) *T. molitor* females after injection of physiological saline (control, red line) or Tenmo-TRP-7 at a concentration 10^-7^ M (blue line) and 10^-5^ M (green line). Additionally, the number of eggs laid was also tested after knockdown of TRP precursor (C). The ds*GmLys* and ds*TRP* was injected into 1-day-old (dsControl 1D, black line and ds*TRP* 1D, violet line) and 4-day-old (dsControl 4D, yellow line and ds*TRP* 4D, pink line) females. The colours in the table indicate an increase (red) in cumulative number of eggs laid relative to the group in the top row of the table. Statistical comparison of cumulative number of laid eggs based on the two-way ANOVA with Tukey’s multiple comparisons test. The values are presented as the means ± SEs. \**p* ≤ 0.05, ns – nonsignificant difference.

### 3.1.4. Changes in larval hatching

Next, it was necessary to examine whether the application of Tenmo-TRP-7 or ds*TRP* affects embryo development and the overall larval hatching rate. Since previous results indicated that TRP can affect the number of eggs laid, determining whether TRP influences later stages of offspring development, such as embryogenesis and hatching, is important. Assessing these parameters allows us to establish whether manipulation of TRP levels affects not only female fertility but also the quality and viability of eggs.

The results revealed that the application of Tenmo-TRP-7 did not significantly affect embryo development or larval hatching success (Fig. 4A and 4B). However, in the case of dsRNA treatment, an increase in egg hatching rate was observed, but only when comparing 1-day-old females injected with ds*TRP* to 4-day-old females injected with either ds*GmLys* (dsControl) or ds*TRP* (Fig. 4C).

**Fig. 4.**
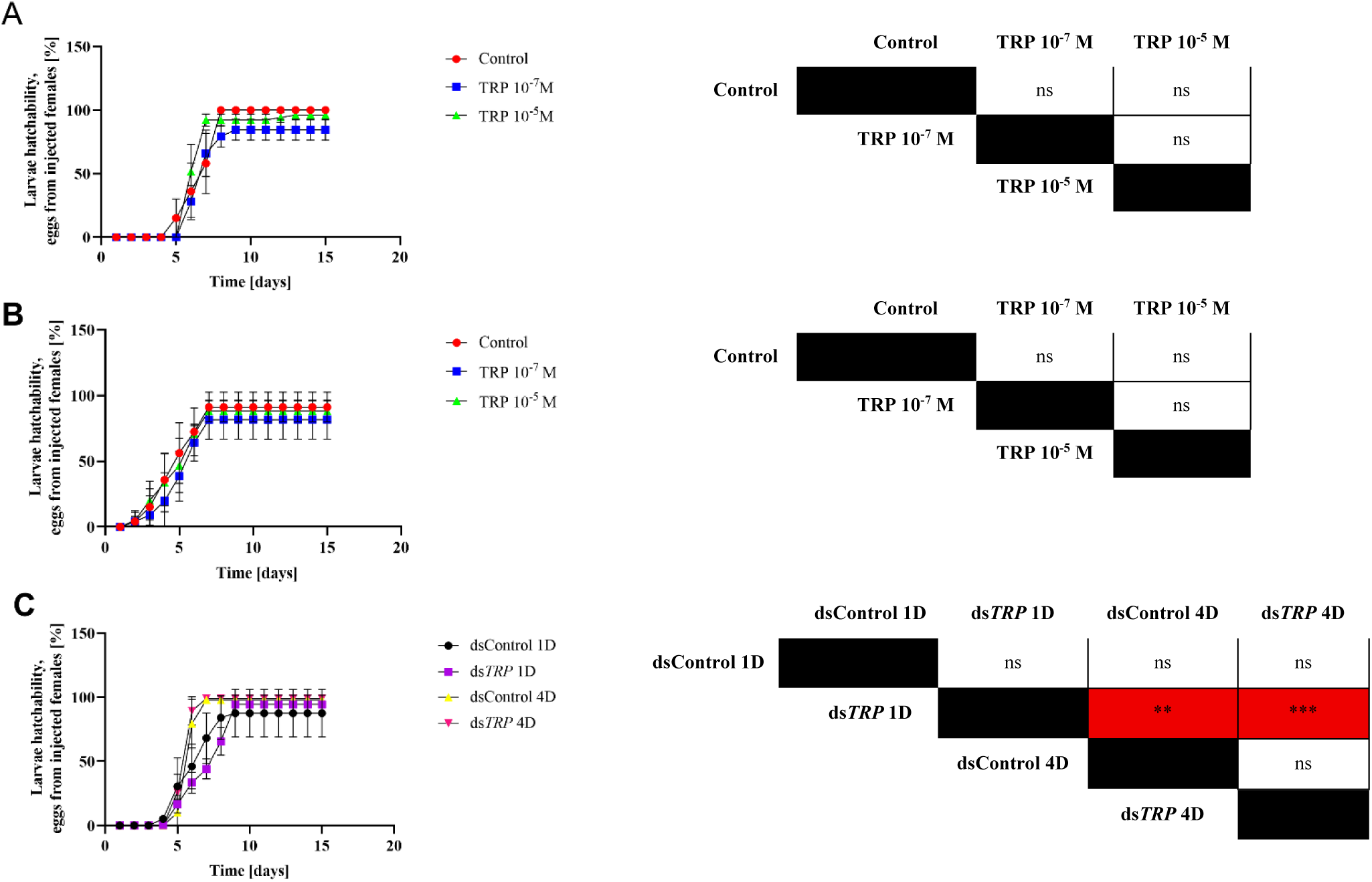
Duration of embryonic development and hatchability of larvae from eggs of 1-day-old (A) and 4-day-old (B) *T. molitor* females after injection of physiological saline (control, red line) or Tenmo-TRP-7 at a concentration 10^-7^ M (blue line) and 10^-5^ M (green line). Additionally, the hatchability of larvae was also tested after knockdown of TRP precursor (C). The dsGmLys and ds*TRP* was injected into 1-day-old (dsControl 1D, black line and ds*TRP* 1D, violet line) and 4-day-old (dsControl 4D, yellow line and ds*TRP* 4D, pink line) females. The colours in the table indicate an increase (red) in cumulative number of eggs laid relative to the group in the top row of the table. The values are presented as the means ± SEs. Statistical comparison of cumulative number of hatchability of larvae based on the two-way ANOVA with Tukey’s multiple comparisons test, \*\**p* ≤0.01, *** *p* ≤0.001, ns – nonsignificant difference.

### 3.1.5. Changes in development of follicular epithelium

To better understand the mechanism underlying the increased number of eggs laid, it was necessary to investigate how Tenmo-TRP-7 application and silencing of the gene encoding its precursor affect the structural organization of the ovary. A particular focus was placed on the permeability of follicular cells that form the ovarian epithelium, as their permeability is essential for transporting nutrients to the developing embryo and may influence both egg quality ^16^. Permeability of follicular cells was expressed as patency index (Fig. 4S) ^19^.

As a first step, changes in epithelial permeability were assessed following the application of the neuropeptide Tenmo-TRP-7. In 1-day-old females 24 hrs after Tenmo-TRP-7 injection, no significant changes were observed (Fig. 5A). However, in 4-day-old females, the administration of Tenmo-TRP-7 led to increased permeability of the follicular epithelium (Fig. 5B). In contrast, dsRNA-based silencing of the *TRP* gene resulted in reduced epithelial permeability in both 1-(Fig. 5C) and 4-day-old females (Fig. 5D).

**Fig. 5.**
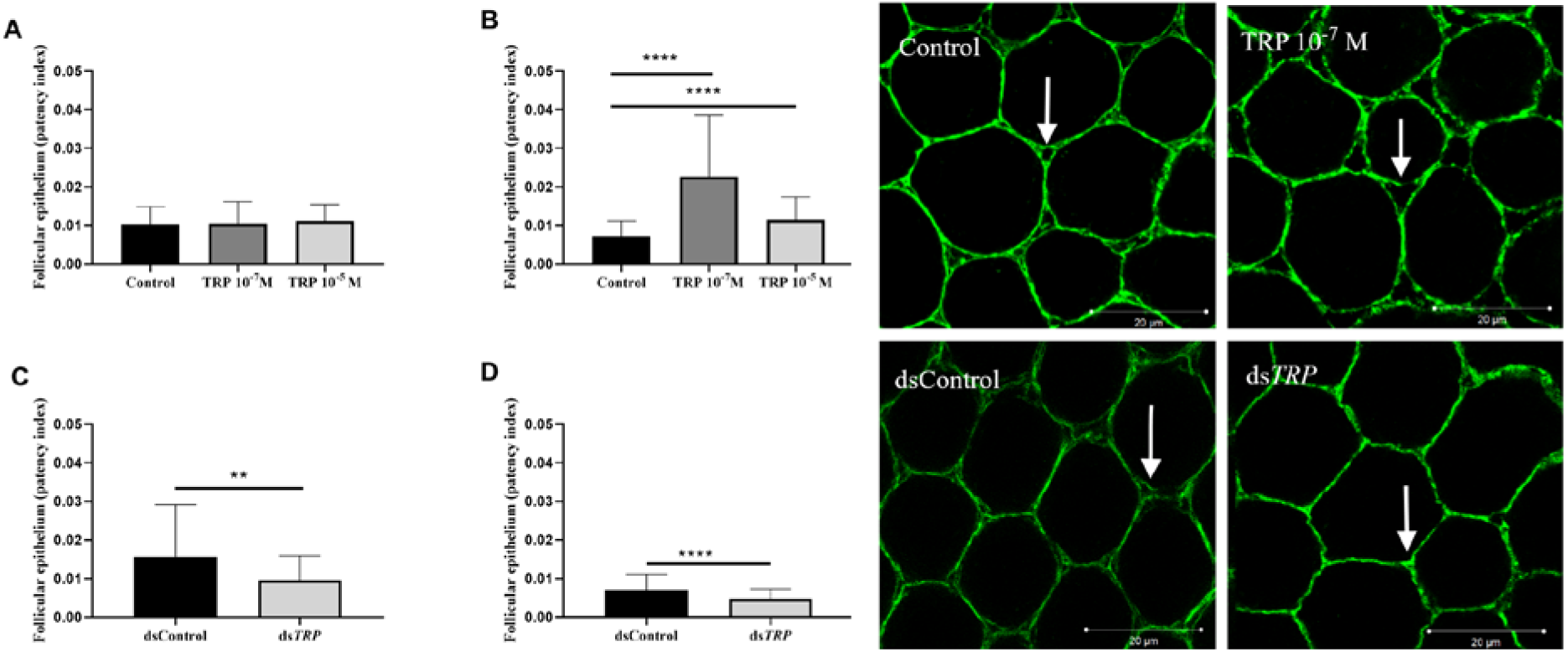
Changes in follicular epithelium permeability (patency index) in 1-day-old (A) and 4-day-old (B) *T. molitor* females after injection of physiological saline (control) or Tenmo-TRP-7 at concentrations of 10^-7^ M and 10^-5^ M. Additionally, the patency index was assessed after ds*GmLys* (dsControl) and ds*TRP* application in 1-day-old (C) and 4-day-old (D) females. Statistical comparison of changes in patency index based on the Kruskal–Wallis test and Mann– Whitney U test. The values are presented as the means ± SDs, \*\**p* ≤0.01, *** *p* ≤0.001, **** *p* ≤0.0001. Micrographs showing the changes in follicular epithelium permeability (patency index) in 4-day-old *T. molitor* females after injection of physiological saline (control), Tenmo-TRP-7 at concentrations of 10^-7^ M, ds*GmLys* (dsControl) or ds*TRP.* The arrows indicate the example of intercellular spaces, whose changes in size affect the permeability of the epithelium. Scale bar – 20 μm.

### 3.1.6. Changes in oocyte maturation

Next, the effect of TRP signalling on the maturation of terminal oocytes was investigated. The terminal oocyte represents the final stage of oocyte development prior to ovulation, and measuring its volume provides insight into whether TRP plays a role in regulating oocyte maturation processes, which are directly linked to female reproductive capacity. Importantly, this stage of oocyte development results from epithelial patency.

In 1-day-old females, Tenmo-TRP-7 application resulted in an increase in terminal oocyte volume at both tested concentrations (Fig. 6A). In contrast, in 4-day-old females, Tenmo-TRP-7 application at a concentration of 10^-5^ M caused a decrease in terminal oocyte volume, whereas no significant changes were observed at 10^-7^ M (Fig. 6B). Knockdown of *TRP* had no effect on oocyte volume in 1-day-old females (Fig. 6C). However, in 4-day-old females, precursor knockdown led to an increase in terminal oocyte volume (Fig. 6D).

**Fig. 6.**
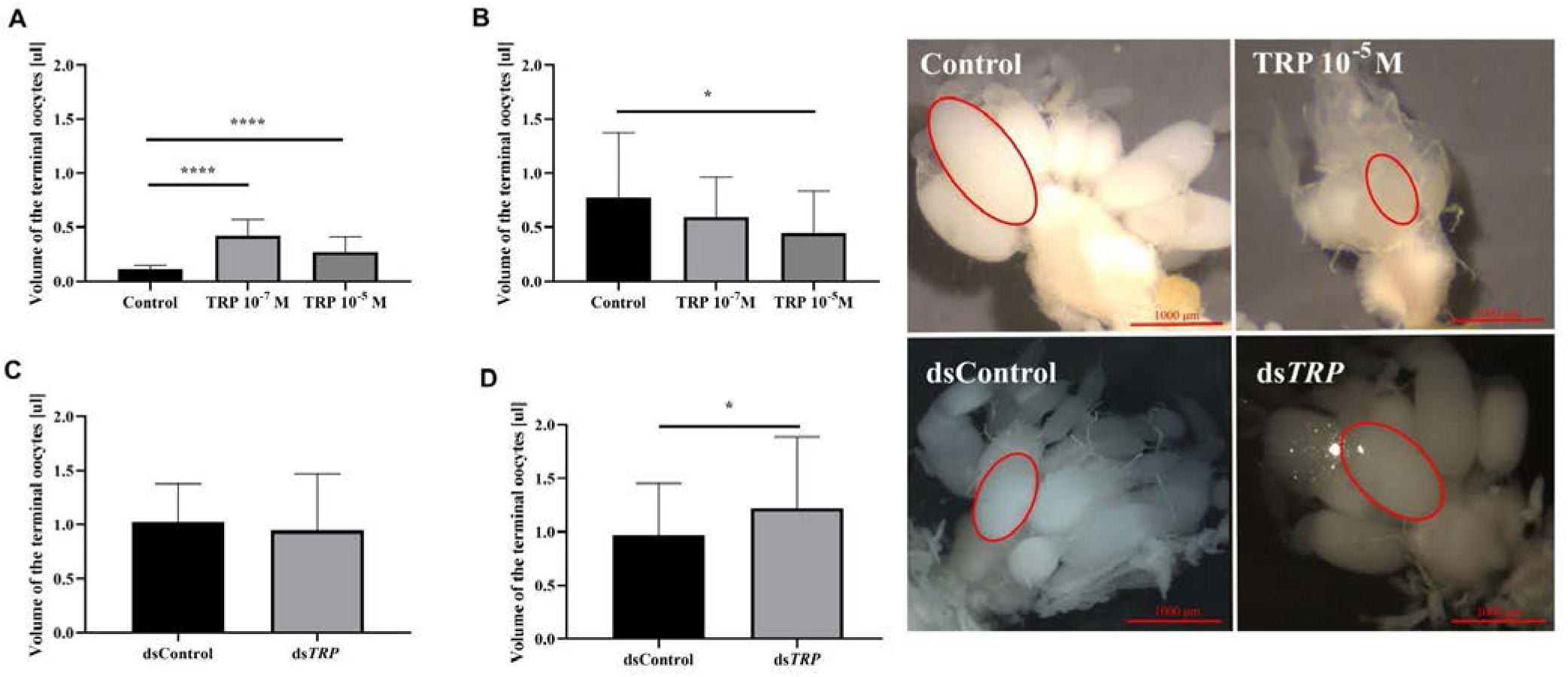
Changes in volume of the terminal oocytes in 1-day-old (A) and 4-day-old (B) *T. molitor* females after injection of physiological saline (control) or Tenmo-TRP-7 at concentrations of 10^-7^ M and 10^-5^ M. Additionally, the volume of terminal oocytes was tested after ds*GmLys* (dsControl) and ds*TRP* application in 1-day-old (C) and 4-day-old (D) females. Statistical comparison of changes in patency index based on the Kruskal–Wallis test and Mann–Whitney U test. The values are presented as the means ± SDs, \**p* ≤ 0.05, **** *p* ≤ 0.0001. Pictures showing the changes in volume of the terminal oocytes in 4-day-old *T. molitor* females after injection of physiological saline (control), Tenmo-TRP-7 at concentrations of 10^-7^ M, ds*GmLys* (dsControl) or ds*TRP.* Red ellipses mark the example of terminal oocyte. Scale bar – 1000 μm.

### 3.1.7. Topical application

Given the established role of TRPs in modulating female reproductive physiology, the potential for applying this knowledge in RNAi-based pest control strategies was examined. Previous research has demonstrated that topically applied dsRNA can penetrate the insect cuticle ^20^. To assess whether dsRNA targeting the *TRP* gene can be delivered via a non-invasive route, a topical application assay was performed. The experiment aimed to evaluate the ability of the dsRNA to penetrate the cuticle of *Tenebrio* females. The dsRNA was applied either with or without the addition of a surfactant (Triton X-100) to females at 1, 4, and 7 days post-emergence. In all the cases, no significant silencing of the *TRP* gene was observed (Fig. 5S). In females that were administered a dsRNA solution with 1% Triton-X and a dsRNA solution with 0.1% Triton-X on the epidermis under the elytra, 100% mortality was observed, which prevented further studies. Due to neuropeptide stability, topical application non-modified Tenmo-TRP-7 was not tested ^21^.

## 4. Discussion

Insects, as the most numerous group of animals in the world, pose a serious threat to global food security by contributing to the destruction of crops ^9^. Moreover, their high nutritional value, short life cycle, and low environmental requirements make them a promising alternative source of, for example, protein for both humans and animals ^22^. However, to fully describe the potential of this species, it is essential to optimize its reproductive cycle, which determines the efficiency of large-scale industrial production. In this study, we demonstrate that Tenmo-TRP-7, as well as the modulation of their signalling pathway using RNAi, may contribute to the regulation of reproductive system activity in *T. molitor* females. This makes them potential candidates for supporting industrial insect rearing and for controlling population.

Our results show a significant positive correlation between expression level of *TRP* and *TRPR* with expression of *Vg* in the fat body. This observation suggests that TRP may stimulate the production of vitellogenin, a key yolk protein synthesized in the fat body ^23^. Up-mentioned finding is consistent with other known functions of this neuropeptide, as TRPs are also involved in regulation of lipid metabolism, gut motility and food intake ^14,15^. In contrast, no significant changes were observed in *Vg* gene expression in the ovaries in response to TRP signaling. However, considering that the fat body is the primary site of vitellogenin synthesis in insects, this lack of ovarian response may reflect tissue-specific regulation. On the other hand, the negative correlation between *TRPR* gene expression in the ovaries and the expression of *TRP* and *TRPR* in the head and fat body may indicate a balancing mechanism or feedback regulation between tissues. The upregulation of *TRP* and *TRPR* genes may be connected to increasing synthesis and releasing of TRP from the nervous system. For this reason observed *TRPR* downregulation in ovaries may be intended to limit excessive stimulation of the TRP pathway in these organs.

The physiological analysis with Tenmo-TRP-7 and ds*TRP* related to the reproduction of *T. molitor* female may indicate the possible role of TRP pathway in regulation of egg production. In 1-day-old females, the administration of Tenmo-TRP-7 at a concentration of 10^-5^ M resulted in a significant increase in *Vg* gene expression in both the fat body and the ovaries. These findings confirm that TRP can directly and/or indirectly stimulate processes related to egg maturation - by activating yolk synthesis in the fat body and potentially enhancing yolk transport and storage in the ovaries. Injection of 4-day-old females with ds*TRP,* just before the peak of reproductive activity resulted in a significant downregulation of *Vg* expression in the fat body but had no effect on *Vg* expression in the ovaries. This can be explained by the fact that the fat body is the primary site of Vg synthesis in insects, and TRP primarily modulates its function ^17,24^. Although Vg detected in the ovaries may partly reflect local synthesis, as shown in other insects such as *Culex quinquefasciatus*, it is generally considered to originate mainly from uptake of fat body-derived protein ^25^. This predominant uptake could explain why no significant expression changes were observed in the ovaries following ds*TRP* treatment. Furthermore, at this stage of the reproductive cycle, oocytes are already supplied with yolk, and their further development likely depends more on processes such as epithelial transport and tissue permeability than on active local synthesis of nutritional components.

Given the observed upregulation of *Vg* expression following Tenmo-TRP-7 and ds*TRP* administration, we investigated whether modulation of TRP signalling also affects the number of eggs laid and their hatching success. The application of synthetic Tenmo-TRP-7 significantly increased the number of eggs laid by young females, and this effect depended on both the age of the individual and the concentration of the administered neuropeptide . In 1-day-old females, the higher Tenmo-TRP-7 concentration (10^-5^ M) was most effective, whereas in 4-day-old females, the lower dose (10^-7^ M) had a greater effect. In contrast, silencing of the gene encoding the *TRP* using dsRNA did not significantly alter the number of eggs laid. This may reflect several non-exclusive compensatory mechanisms. First, the neuroendocrine regulation of insect reproduction involves overlapping signaling pathways; other neuropeptides such as allatotropin, bursicon, or short neuropeptide F may partially compensate for the absence of TRP, maintaining a baseline level of oviposition ^26,27^. Second, the RNAi effects were assessed eight days post-injection, by which time the first reproductive cycle had already been completed and the ovary had entered a more asynchronous state, potentially masking earlier changes in reproductive dynamics ^19^. Moreover, already synthesized TRPs or their reservoirs in peripheral tissues may have sustained ovarian activity during this period. Due to the nature of neuropeptides, which are often stored in secretory granules and released in a regulated manner rather than being continuously synthesized, their effects can persist even when transcriptional activity is transient or reduced ^28^. Finally, it is also possible that TRPs act more as reproductive facilitators than essential components - their presence enhances egg-laying efficiency, but is not strictly required for the process to occur. From the perspective of insect farming for food, an important aspect was also the assessment of the effect of TRP on embryo development and hatching success. The data revealed that neither the administration of synthetic TRP nor its silencing had any impact on embryogenesis or larval survival. This finding indicates that TRPs are not teratogenic and do not negatively affect offspring quality, which is crucial in the context of industrial insect production - it allows for the stimulation of reproduction without the risk of compromising the quality of the resulting larvae.

The ability of follicular epithelium to transport nutrients into the oocyte is crucial for both the quality and quantity of eggs laid ^29^. Further analyses revealed that TRPs also affect the permeability of the ovarian epithelium, which plays an important role in the transport of nutrients to developing oocytes ^29^. In 4-day-old females, Tenmo-TRP-7 administration increased ovarian patency index, which may enhance the efficiency of nutrient delivery to eggs, leading to faster accumulation of yolk reserves and thereby shortening oocyte growth time. In contrast, *TRP* gene silencing reduces this parameter regardless of age. This partially explains the previously observed differences in egg numbers and may be significant not only for increasing production under rearing conditions but also as a potential target for novel plant protection strategies. Although silencing of *TRP* expression alone did not reduce the number of eggs laid, the observed decrease in ovarian patency index suggests that more effective or sustained disruption of TRP signaling could impair nutrient transport to developing oocytes. Due to the nature of neuropeptides, which are often stored in vesicles and released in a regulated manner, transient silencing may result in only limited effects. In turn, stronger or prolonged inhibition could eventually reduce reproductive output, offering a promising avenue for pest control that does not rely on conventional insecticidal agents.

Building on these observations, we next assessed the impact of Tenmo-TRP-7 on terminal oocyte volume at different female ages. Analysis of terminal oocyte volume revealed that in 1-day-old females, Tenmo-TRP-7 administration increased terminal oocyte volume, suggesting a stimulatory role of TRP in early oocyte maturation. In contrast, in 4-day-old females, administration of a high TRP concentration (10^-5^ M) led to a reduction in oocyte volume. Given the known myostimulatory properties of TRPs, this effect may result from accelerated oocyte passage through the ovarioles and oviducts ^24^. Faster transport may reduce the time available for vitellogenin accumulation, resulting in smaller oocytes but an increased number of eggs laid. Interestingly, *TRP* silencing in older females had the opposite effect, leading to an increase in terminal oocyte volume. This may reflect a compensatory mechanism whereby delayed oocyte transport prolongs exposure to circulating Vg, allowing for greater resource accumulation and ultimately larger oocyte size, despite lower permeability of follicular epithelium.

The separate but very important issue is potential usage of obtained results in pest control and insect mass rearing. Research by Killiny et al. ^30^ show that dsRNA may penetrate cuticle of *Diaphorina citri*. Our experiment involving the application of dsRNA showed that ds*TRP* probably is unable to penetrate the cuticular barrier of *T. molitor*. Similar results were reported by Finetti et al. ^20^ in studies on *Rhodnius prolixus* (Hemiptera), where topically applied dsRNA had no biological effect and did not reduce gene expression. The lack of efficacy was likely due to the presence of a waxy layer on the surface of the abdomen, which protects the body from the penetration of hydrophilic molecules such as dsRNA ^20^. Beetles (Coleoptera), including *T. molitor*, possess a highly developed, hydrophobic, and often more strongly sclerotized cuticle with a lipid layer, which may protect against the penetration of large, hydrophilic molecules like dsRNA even more effectively than in Hemiptera ^31^. Triton X-100 may be insufficient to overcome this barrier. Numerous studies have shown that RNAi efficacy varies significantly across insect orders - Diptera and Lepidoptera are often resistant, while in Coleoptera and Hemiptera, the efficiency is highly variable ^32,33^. Therefore, the next step will require the use of nanocarriers such as chitosan or liposomes, as well as the implementation of alternative delivery methods, for example, administration through feeding. In addition, we did not employ non-modified neuropeptides in our experiments. This decision was based on the fact that non-modified neuropeptides are characterized by a short half-life, which substantially limits their biological activity ^12^. Moreover, they often exhibit limited receptor specificity and are rapidly removed from the organism ^34^. Their synthesis and storage are also associated with high costs and considerable technical challenges ^35^.

In the discussion concerning the role of the TRP system on beetle reproduction we must mention another insect neuropeptide family, natalisins (NTL). Notably, Tenmo-TRP-7 shares functional similarities with NTL, which likely evolved from the same ancestral gene and exhibits a similar C-terminal motif (e.g., FxxxRa). Despite acting through distinct GPCRs, TRP and NTL may have overlapping roles in reproductive regulation. Studies in *Tribolium castaneum* and *Drosophila melanogaster* indicate that NTL is essential for normal mating behavior and egg laying, and its silencing significantly reduces fecundity without affecting embryonic development - an effect comparable to the non-teratogenic action of TRP in *T. molitor* ^36^. However, the use of specific dsRNA targeting TRP often produced physiological effects opposite to those observed following the injection of Tenmo-TRP-7, suggesting a direct role of TRP signaling in *T. molitor* reproduction.

In summary, TRP system exerts a multi-level influence on the reproduction of *Tenebrio molitor* females, affecting processes ranging from the activation of Vg synthesis and regulation of ovarian epithelial permeability to the modulation of egg number. Their use has the potential to enhance rearing efficiency without negatively impacting offspring development, making them attractive tools for improving insect production for feed and food applications. At the same time, targeting the same signaling pathway in pest species could form the basis for the development of modern, selective, and environmentally safe plant protection strategies.

## 5. Materials and methods

### 5.1. Insects

The model organisms used in the study were female *T. molitor* beetles. Insects were reared at the Department of Animal Physiology and Developmental Biology (Adam Mickiewicz University, Poznań, Poland) in MIR 154-PE incubators (PHCbi, Singapore, Republic of Singapore) in full dark conditions at 28°C and 65 ± 5% humidity. Adult insects were fed flour and apples. Males were added to each container with females to ensure adequate ovarian growth and development. Both sexes were kept in one container at a 2:1 ratio (two females per one male, 15 insects per box), as the presence of five males increased reproductive success ^37^. The ages of the females used for specific experiments are described in the sections below.

### 5.2. Tested compounds

#### 5.2.1. Tenmo-TRP-7

The neuropeptide Tenmo-TRP-7 (amino acid sequence: MPRQSGFFGMRa) synthesized by Creative Peptides (Shirley, NY, USA; purity >95% HPLC) was used in the experiment. The effect of this neuropeptide on key reproductive parameters were assessed 24 hours after injection at two concentrations: 10⁻⁵ M and 10⁻⁷ M in physiological saline (PS).

#### 5.2.2. dsRNA directed against the TRP precursor (ds*TRP*)

Double-stranded RNA (dsRNA) targeting the *TRP* was synthesized using a modified protocol based on the methods described by Konopińska et al ^38^. The first step was isolation of total RNA from adult *T. molitor* and *Galleria mellonella* larvae (used as a control of dsRNA treatment) using the Quick-RNA Mini-Prep Kit (Zymo Research, Irvine, CA, USA). Due to its lack of specificity towards *Tenebrio* genes, dsRNA targeting *G. mellonella* lysozyme does not induce RNAi and does not affect the biological parameters studied. The extracted RNA was treated with the TurboDNA-free Kit (Zymo Research, Irvine, CA, USA) to eliminate genomic DNA contamination. RNA concentration and purity were assessed using a DS-11 spectrophotometer (DeNovix, Inc., Wilmington, DE, USA). Complementary DNA was synthesized from the purified RNA using the LunaScript® RT SuperMix Kit (New England Biolabs, Ipswich, MA, USA), with a no-RT control included to verify the absence of DNA contamination. PCR amplification of specific gene fragments was performed using the KAPA2G Fast ReadyMix PCR Kit (KAPA Biosystems, Sigma‒Aldrich, St. Louis, MO, USA), with cDNA serving as the template. The primer sequences used are listed in Table 4. To verify product integrity, the PCR amplicons were analyzed by electrophoresis on a 2% TAE agarose gel stained with ethidium bromide. The PCR products were then purified using the PCR/DNA Clean-Up Kit (EURx, Poland). dsRNA was synthesized using the HighYield T7 RNA Synthesis Kit (Jena Bioscience GmbH, Jena, Germany) following the manufacturer’s instructions, with the transcription step extended to 4 hours at 37°C. The residual DNA was further removed by an additional TurboDNA-free (Thermo Fisher Scientific, Waltham, MA, USA) treatment. For annealing, samples were heated to 95°C and allowed to cool gradually overnight in a thermocycler. Obtained dsRNA was purified through wash using 5 M ammonium acetate (NH₄OAc) (Thermo Fisher Scientific, Waltham, MA, USA) and then gradient wash with molecular-grade ethanol (99% to 70%). The dsRNA pellet was dissolved in nuclease-free water and stored at −20°C until use. Gene silencing was achieved by injecting adult beetles with 2 µg of dsRNA (1 µg/µL in 2 µL of nuclease-free water).

Prior to the main experiments, the knockdown efficiency was assessed by extracting total RNA (*n* = 3, pools of three females per time point) at various intervals post-injection. The samples were processed as described above, and relative gene expression was quantified by RT-qPCR (as detailed in Section 4.4.). On the basis of these results, day 8 post-injection was selected as the optimal time point, showing a clear reduction in *TRP* gene expression (Fig. 7).

**Fig. 7.**
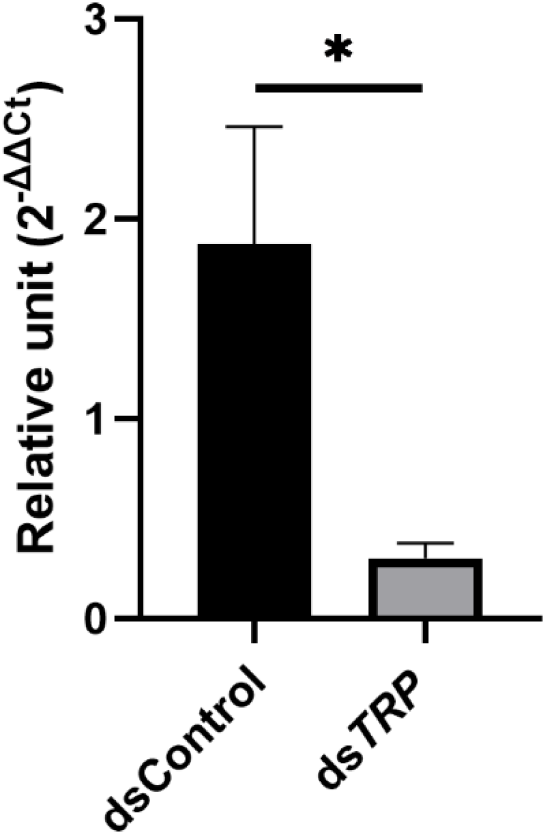
Efficiency of TRP knockdown on the 8^th^ day after dsRNA injection. dsControl – The mRNA quantity of TRP was measured in relation to *Galeria melonella* lysozyme (GmLys) as a negative control by RT-qPCR. The values are presented as the means ± SDs, \**p* ≤ 0.05.

### 5.3. Injection and sample collection

Insects were anaesthetized for 7 minutes using endogenous carbon dioxide, followed by surface sterilization in ethanol and rinsing with distilled water. Then, with a microliter syringe (Hamilton Company, Reno, NV, USA), 2 µL of the physiological saline solution (control) or Tenmo-TRP-7 at concentrations of 10^-7^ M and 10^-5^ M was injected under the coxa of the third pair of legs. Additionally for the RNAi experiment females were injected with 2 µL of ds*GmLys* (dsControl) or ds*TRP.* Since *T. molitor* commonly experiences injuries and the injection of sterile saline has minimal impact, the study did not include an additional control group of unmanipulated females ^39^.

### 5.4. Expression levels of TRP, TRPR and vitellogenin in 1-7-day-old females of *Tenebrio molitor*

The head, fat body and ovaries were dissected from noninjected females under sterile conditions via a Zeiss Stemi 504 microscope (Zeiss, Jena, Germany) between day 1 and day 7 after eclosion. The dissected tissues were placed into individual Eppendorf tubes containing 300 µL of lysis buffer (Zymo Research, Irvine, CA, USA). The samples were immediately flash-frozen in liquid nitrogen and stored at −80 °C until further processing. Prior to RNA extraction, the tissues were homogenized for 1 minute using a pellet homogenizer (Kimble Chase, Vineland, NJ, USA). RNA was then extracted using the Quick-RNA Mini-Prep Kit (Zymo Research, Irvine, CA, USA) following the manufacturer’s instructions. To remove any contaminating DNA, the samples were treated with the Turbo DNA-free Kit (Thermo Fisher Scientific, Waltham, MA, USA). RNA concentration and purity were assessed using a DS-11 spectrophotometer (DeNovix, Inc., Wilmington, DE, USA).

cDNA was synthesized from equal concentrations of RNA templates (200 ng) via a LunaScript® RT SuperMix Kit (New England Biolabs, Ipswich, MA, USA). Additional quality control (no RT control) was also performed. For RT‒qPCR analysis, the expression levels of genes encoding TRP precursor and TRPR were analysed. Moreover, in both experiments, the expression level of *Vg* was tested. The primers used in the experiment were synthesized by the Institute of Biochemistry and Biophysics of the Polish Academy of Science (Warsaw, Poland). Moreover, to confirm our results, the amplicons were sequenced by the Molecular Biology Techniques Laboratory (Faculty of Biology, Adam Mickiewicz University, Poznań) and compared with data available in a public database (https://www.ncbi.nlm.nih.gov). The expression of selected genes was normalized on the basis of the expression level of the gene encoding *T. molitor* ribosomal protein L13a (*TmRpL13a*) ^40^. The sequences of the primers used are available in Table 1. The efficiency of the primers used ranged from 92-108%. RT‒qPCR analysis was performed on a QuantStudio™ 3 Real-Time PCR System, 96-well, 0.2 mL (Applied Biosystems™, Waltham, MA, USA). In one biological replication, tissues from five individuals (for the ovaries) or 10 individuals (head or fat body) were collected for each sample. The relative expression was calculated via the method described by Pfaffl ^41^.

**Tab. 1.**
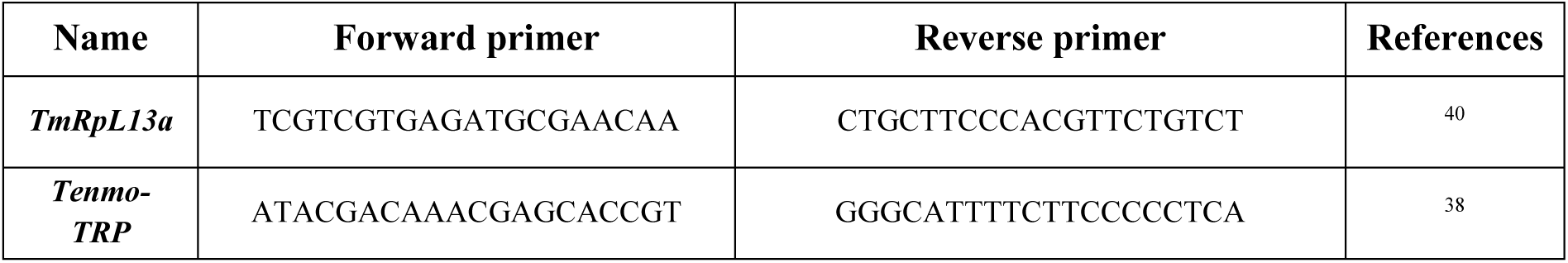
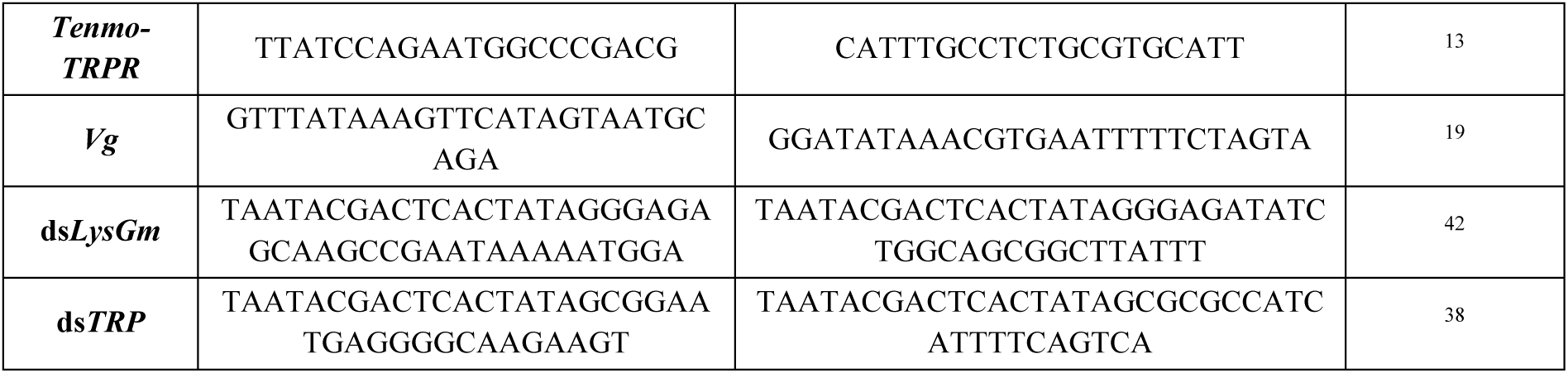
Sequences of the primers used in the PCR analysis.

### 5.5. Expression levels of gene encoding vitellogenin after the application of Tenmo-TRP-7 and dsRNA

Female *Tenebrio* were injected on the 1st and 4th day after eclosion. After the application of Tenmo-TRP-7 (24 hours) or dsRNA (8 days), the ovaries and fat body were isolated from each individual. The samples were prepared and processed as described in section 4.4.

### 5.6. Survival experiment

Female *T. molitor* at 1 and 4 day post-eclosion were injected with the tested solutions. Survival was recorded daily at the same time for 15 consecutive days. Five individuals kept in a plastic box were considered one biological replicate, and each experimental variant was performed in five replicates.

### 5.7. Changes in egg production

To evaluate the impact of Tenmo-TRP-7 on oviposition, females were injected on days 1 and 4, after eclosion. For the RNAi experiment 1- and 4-day-old females were injected and the eggs were collected on the 8th day post-injection. Egg laying was monitored over the following 10 days. Eggs were collected daily using the method described by Rosiński et al. ^43^ in patent PL No. 217868B1, which involved sieving the substrate through a mesh with 1.15 mm openings. After each collection, the culture medium was replaced. Each experimental group consisted of five biological replicates, with five females per replicate. The cumulative number of eggs laid per female was calculated for each group.

### 5.8. Changes in larval hatching

Changes in egg hatchability were examined based on the method described by Walkowiak Nowicka et al. ^19,44^. Female *T. molitor* at 1 and 4 days post-eclosion were injected with Tenmo-TRP-7 at concentrations of 10^-7^ M and 10^-5^ M. Eggs were collected the 4th day after injection. For the RNAi experiment 1 and 4-day-old females were injected, and eggs were collected on the 8^th^ day. Subsequently, the collected eggs were placed in Petri dishes with a small amount of coarse grain flour and kept in an incubator at a constant temperature of 28°C and 65 ± 5% relative humidity. Larval hatchability was monitored for the next 10 days. Every day, the hatched larvae were removed to prevent them from eating the eggs. Five replicates were performed for each variant. Larval hatchability was calculated as the percentage of larvae hatched relative to the total number of eggs laid.

### 5.9. Changes in follicular epithelium development

The effects of Tenmo-TRP-7 application on the development of the follicular epithelium were assessed according to the method described by Walkowiak-Nowicka et al. ^44^. The experiment was conducted on 1- and 4-day-old females. Ovaries were isolated 24 hours after injection of Tenmo-TRP-7 at concentrations of 10^-7^ M and 10^-5^ M or 8 days after infection with ds*GmLys* or ds*TRP*, and fixed in 4% paraformaldehyde solution in PS. Samples were then incubated with 0.1% Triton X-100 (Merck Life Science, Poland). To visualize the F-actin cytoskeleton in follicular epithelial cells, Oregon Green® 488 phalloidin (Thermo Fisher Scientific, Waltham, MA, USA) was used in a solution of 1% bovine serum albumin (BSA prepared in PSA) and incubated for 20 minutes at room temperature in the dark. The ovaries were then transferred onto microscope slides and secured using a mixture of glycerol, phosphate-buffered saline (PBS), and DABCO. Follicular epithelium development was evaluated using a Carl Zeiss LSM 510 confocal microscope operated with the AxioVision SE64 Rel. 4.9.1 software. The patency index was calculated using the following formula: P = S₄ / (S₁ + S₂ + S₃), where S₄ is the surface area of the intercellular space, calculated as the average area of all visible intercellular spaces in a given experimental variant; S₁, S₂, and S₃ are the surface areas of follicular cells adjacent to a given intercellular space, calculated as the average area of these cells from the microscope image (Fig. 4S).

Each experimental variant included 10 biological replicates. For each ovary, 3 random images were taken and analysed.

### 5.10. Changes in oocyte maturation

Changes in the volume of terminal oocytes after Tenmo-TRP-7 application were examined using the method described by Walkowiak-Nowicka et al. ^19^ and Pszczolkowski et al ^45^. 1- and 4r-day-old females were injected with TRP at concentrations of 10⁻⁵ M and 10⁻⁷ M, and after 24 hours the ovaries were isolated. After the application of ds*GmLys* or ds*TRP*, the isolation was performed on the 8^th^ day post-injection. Next samples were fixed in a 4% paraformaldehyde solution in physiological saline. Photographs were then taken using an Olympus SZX12 stereomicroscope with Olympus DP-Soft Analysis software version 3.0. For the analysis, the length and width of terminal oocytes were measured. The oocyte volume was calculated using the formula: V = 4/3 × π × a × r². Ten biological replicates were performed for each experimental variant.

### 5.11. Topical application

To assess the ability of dsRNA targeting the *TRP* to penetrate the insect cuticle, a topical application approach was employed. The experiment was conducted with several variants.

In the first experimental variant, 1-day-old insects, whose cuticle had not yet fully melanized, and 7-day-old adult individuals were used. Both groups were divided into two subgroups. In the first group 2 µL of dsRNA was applied under the elytra. The second group had 2 µL of dsRNA applied to the intersegmental membrane. For the second experimental variant, a mixture of dsRNA with a 1% Triton X-100 solution was used. 1-day-old and 7-day-old adults were treated with 2 µL of this mixture applied under the elytra. The third variant involved the use of a 0.1% Triton X-100 solution mixed with dsRNA, which was applied under the elytra and intersegmental membrane of one and 7-day-old adults. Additionally, to confirm the efficacy of dsRNA, a positive control was performed by injecting dsRNA directly into the insect hemolymph.

Due to low stability of neuropeptides in the environment, Tenmo-TRP-7 topical application has not been tested.

### 5.12. Statistical analysis

Statistical analyses were carried out using GraphPad Prism 9 (licensed to Adam Mickiewicz University). Survival data were analyzed using Kaplan–Meier survival curves, and differences between curves were assessed using the log-rank (Mantel–Cox) test. Correlation analyses were performed using the Pearson correlation coefficient method via the SRplot platform (https://www.bioinformatics.com.cn/srplot). For data following a normal distribution, one-way ANOVA or two-way ANOVA by Tukey’s post hoc test was applied. In cases where data did not meet normality assumptions, the Kruskal–Wallis test with Dunn’s post hoc test or Mann– Whitney U test was used.

## Acknowledgement

This research was supported by Grant No. 2021/41/B/NZ9/01054 from the National Science Centre (Poland). We would also like to thank Natalia Bylewska, Radosław Gmyrek, Karolina Kozłowska, and Sara Tchórzewska for their help in rearing *T. molitor*.

During the preparation of this work, the authors used Rubriq software (American Journal Experts (AJE)) for language correction. After using this tool, the authors reviewed and edited the content as needed and took full responsibility for the content of the publication.

## Author contributions

Conceptualization - NK, KWN and AU; Sample collection – NK, KWN and SC; Methodology – NK, KWN and GN; Performing analyses – NK and KWN, Visualization – NK and KWN; Writing – Original Draft Preparation – NK, Writing – Review & Editing – NK, KWN, SC, GN, and AU; Supervision – KWN and AU.

## References

1. Gerland, P., et al. World Population Prospects 2022: Summary of Results. (2022).

2. Van Dijk, M., Morley, T., Rau, M. L. & Saghai, Y. A meta-analysis of projected global food demand and population at risk of hunger for the period 2010–2050. Nat Food 2, 494–501 (2021).

3. Steinfeld, H., Gerber, P., Wassenaar, T. D., Castel, V. & De Haan, C. Livestock’s Long Shadow: Environmental Issues and Options. (Food & Agriculture Org., 2006).

4. EFSA Panel on Nutrition, N. F., et al. Safety of dried yellow mealworm (*Tenebrio molitor* larva) as a novel food pursuant to Regulation (EU) 2015/2283. EFSA Journal 19, e06343 (2021).

5. Moruzzo, R. et al. Mealworm (*Tenebrio molitor*): Potential and challenges to promote circular economy. Animals 11, 2568 (2021).

6. Kotsou, K., Chatzimitakos, T., Athanasiadis, V., Bozinou, E. & Lalas, S. I. Exploiting agri-food waste as feed for *Tenebrio molitor* larvae rearing: A review. Foods 13, 1027 (2024).

7. Oonincx, D. G. A. B. & De Boer, I. J. M. Environmental impact of the production of mealworms as a protein source for humans–a life cycle assessment. PLoS One 7, e51145 (2012).

8. Zahra, A. A., Bedewy, M. M. M., Elsaffany, A. H., Metwaly, K. H. & Gad, H. A. Insecticidal effect of ozone on larvae of *Tenebrio molitor* and their enzyme activity. Journal of Plant Diseases and Protection 132, 124 (2025).

9. Spochacz, M., Chowański, S., Walkowiak-Nowicka, K., Szymczak, M. & Adamski, Z. Plant-derived substances used against beetles–pests of stored crops and food–and their mode of action: A review. Compr Rev Food Sci Food Saf 17, 1339–1366 (2018).

10. Jayaprakas, C. A., Tom, J. & Sreejith, S. Impact of insecticides on man and environment. in Biomedical applications and toxicity of nanomaterials 751–768 (Springer, 2023).

11. https://www.fao.org/faostat/en/#data/.

12. Van Hiel, M. B. et al. Neuropeptide receptors as possible targets for development of insect pest control agents. Neuropeptide systems as targets for parasite and pest control 211–226 (2010).

13. Urbański, A. et al. A possible role of tachykinin-related peptide on an immune system activity of mealworm beetle, *Tenebrio molitor* L. Dev Comp Immunol 120, 104065 (2021).

14. Toprak, U. The role of peptide hormones in insect lipid metabolism. Front Physiol 11, 434 (2020).

15. Ahrentløv, N. et al. Protein-responsive gut hormone tachykinin directs food choice and impacts lifespan. Nat Metab 1–23 (2025).

16. Gerber, G. H. Reproductive behaviour and physiology of *Tenebrio molitor* (coleoptera: Tenebrionidae): II. Egg development and oviposition in young females and the effects of mating. Can Entomol 107, 551–559 (1975).

17. Kodrik, D., Frydrychová, R. Č., Hlavkova, D., Habuštová, O. S. & Štěrbová, H. Unusual functions of insect vitellogenins: minireview. Physiol Res 72, S475 (2023).

18. Gong, W. et al. Transcriptome and Neuroendocrinome Responses to Environmental Stress in the Model and Pest Insect *Spodoptera frugiperda*. Int J Mol Sci 26, (2025).

19. Walkowiak-Nowicka, K. et al. Antheraea peptide and its analog: Their influence on the maturation of the reproductive system, embryogenesis, and early larval development in *Tenebrio molitor* L. beetle. PLoS One 17, e0278473 (2022).

20. Finetti, L., Benetti, L., Leyria, J., Civolani, S. & Bernacchia, G. Topical delivery of dsRNA in two hemipteran species: evaluation of RNAi specificity and non-target effects. Pestic Biochem Physiol 189, 105295 (2023).

21. Caers, J. et al. More than two decades of research on insect neuropeptide GPCRs: an overview. Front Endocrinol (Lausanne) 3, 151 (2012).

22. Huis, A. van, et al. Edible insects: future prospects for food and feed security. (2013).

23. Wu, Z., Yang, L., He, Q. & Zhou, S. Regulatory mechanisms of vitellogenesis in insects. Front Cell Dev Biol 8, 593613 (2021).

24. Nässel, D. R. Tachykinin-related peptides in invertebrates: a review. Peptides (N.Y.) 20, 141–158 (1999).

25. Moura, A. S., Costa-da-Silva, A. L., Peixoto, P. S., Maciel, C. & Cardoso, A. F. Vitellogenin genes are transcribed in *Culex quinquefasciatus* ovary. Mem Inst Oswaldo Cruz 118, e220143 (2023).

26. Konopińska, N. et al. The allatotropin/orexin system as an example of immunomodulatory properties of neuropeptides. Insect Biochem Mol Biol 171, 104149 (2024).

27. Fujinaga, D., Shiomi, K., Yagi, Y., Kataoka, H. & Mizoguchi, A. An insulin-like growth factor-like peptide promotes ovarian development in the silkmoth *Bombyx mori*. Sci Rep 9, 18446 (2019).

28. Nässel, D. R. Neuropeptides in the nervous system of *Drosophila* and other insects: multiple roles as neuromodulators and neurohormones. Prog Neurobiol 68, 1–84 (2002).

29. Row, S., Huang, Y.-C. & Deng, W.-M. Developmental regulation of oocyte lipid intake through ‘patent’follicular epithelium in *Drosophila melanogaster*. iScience 24, (2021).

30. Killiny, N., Hajeri, S., Tiwari, S., Gowda, S. & Stelinski, L. L. Double-stranded RNA uptake through topical application, mediates silencing of five CYP4 genes and suppresses insecticide resistance in *Diaphorina citri*. PLoS One 9, e110536 (2014).

31. Zhu, K. Y. & Palli, S. R. Mechanisms, applications, and challenges of insect RNA interference. Annu Rev Entomol 65, 293–311 (2020).

32. Terenius, O. et al. RNA interference in Lepidoptera: an overview of successful and unsuccessful studies and implications for experimental design. J Insect Physiol 57, 231–245 (2011).

33. Singh, I. K., Singh, S., Mogilicherla, K., Shukla, J. N. & Palli, S. R. Comparative analysis of double-stranded RNA degradation and processing in insects. Sci Rep 7, 17059 (2017).

34. Scherkenbeck, J. & Zdobinsky, T. Insect neuropeptides: structures, chemical modifications and potential for insect control. Bioorg Med Chem 17, 4071–4084 (2009).

35. Shi, M. & McHugh, K. J. Strategies for overcoming protein and peptide instability in biodegradable drug delivery systems. Adv Drug Deliv Rev 199, 114904 (2023).

36. Jiang, H. et al. Ligand selectivity in tachykinin and natalisin neuropeptidergic systems of the honey bee parasitic mite *Varroa destructor*. Sci Rep 6, 19547 (2016).

37. Worden, B. D. & Parker, P. G. Polyandry in grain beetles, *Tenebrio molitor*, leads to greater reproductive success: material or genetic benefits? Behavioral Ecology 12, 761–767 (2001).

38. Konopińska Natalia et al. Tachykinin-related peptide signalling is important for the immune response of the mealworm beetle *Tenebrio molitor* L. – in review

39. Goerlinger, A., Develay, C., Balourdet, A., Rigaud, T. & Moret, Y. Infection risk by oral contamination does not induce immune priming in the mealworm beetle (*Tenebrio molitor*) but triggers behavioral and physiological responses. Front Immunol 15, 1354046 (2024).

40. Jacobs, C. G. C. et al. Endogenous egg immune defenses in the yellow mealworm beetle (*Tenebrio molitor*). Dev Comp Immunol 70, 1–8 (2017).

41. Pfaffl, M. W. A new mathematical model for relative quantification in real-time RT–PCR. Nucleic Acids Res 29, e45–e45 (2001).

42. Zanchi, C., Johnston, P. R. & Rolff, J. Evolution of defence cocktails: Antimicrobial peptide combinations reduce mortality and persistent infection. Mol Ecol 26, 5334–5343 (2017).

43. Rosiński, G., Łukowski, A. & Czarniewska, E. PL Patent No 217868 B1 The method of obtaining cleaned Tenebrio molitor eggs and their application. Polish: Sposób pozyskiwania czystych jaj chrząszcza Tenebrio molitor oraz ich zastosowanie. Polish Patent Office (2011).

44. Walkowiak-Nowicka, K., Mirek, J., Chowański, S., Sobkowiak, R. & Słocińska, M. Plant secondary metabolites as potential bioinsecticides? Study of the effects of plant-derived volatile organic compounds on the reproduction and behaviour of the pest beetle *Tenebrio molitor*. Ecotoxicol. Environ. Saf 257, 10–1016 (2023).

45. Pszczolkowski, M. A., Peterson, A., Srinivasan, A. & Ramaswamy, S. B. Pharmacological analysis of ovarial patency in *Heliothis virescens*. J Insect Physiol 51, 445–453 (2005).

