## Supplementary materials for "Effects of tachykinin-related peptides on the reproductive system of *Tenebrio molitor* females: implications for insect breeding and pest control"

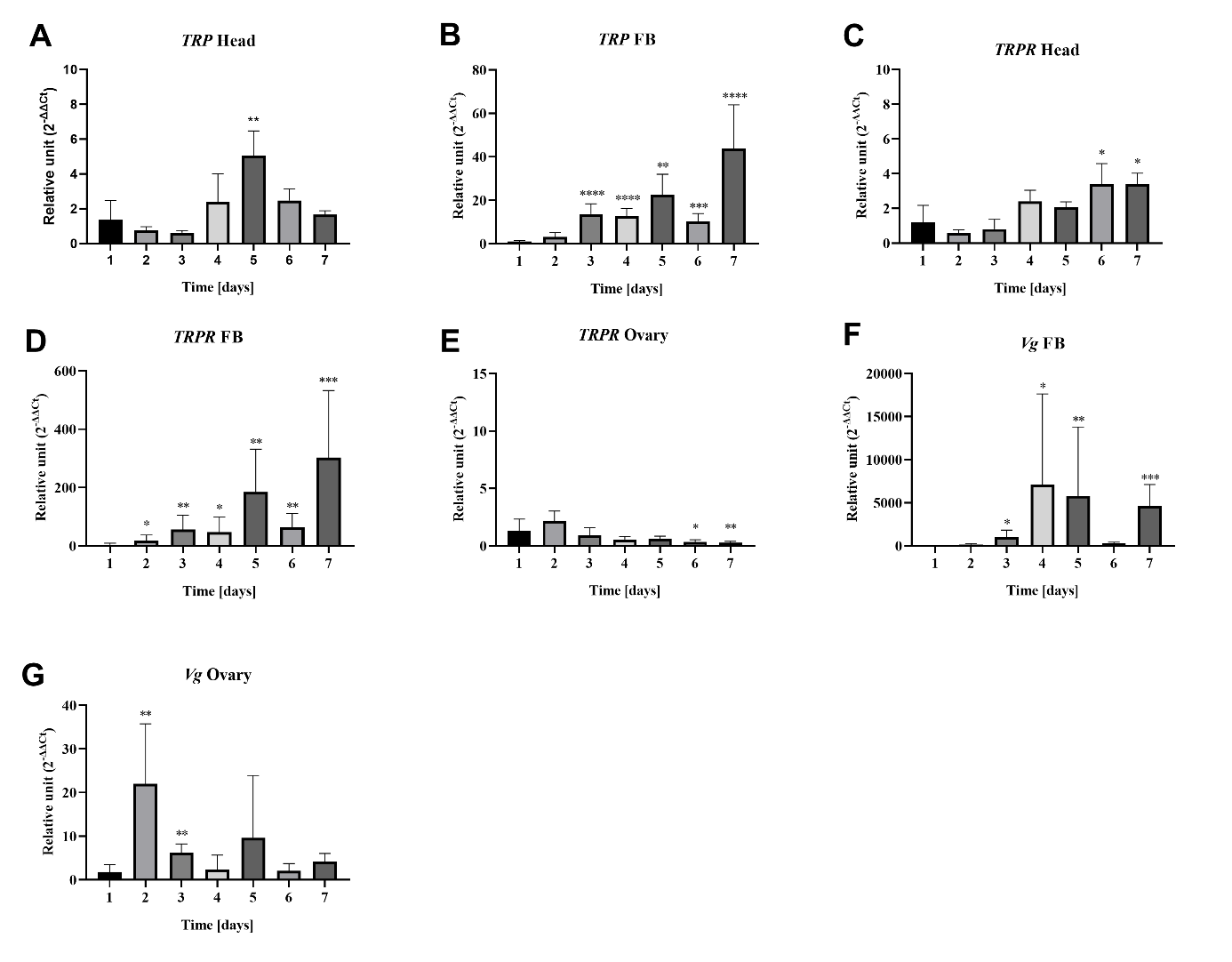

**Fig. 1S**. Changes in the expression levels of genes encoding TRP precursor in the head (A) and fat body (B), TRP receptor (TRPR) in the head (C), fat body (D), and ovaries (E), and genes encoding vitellogenin (Vg) in the fat body (F) and ovaries (G) of 1–7-day-old *T. molitor* females. Values are presented as means ± SD. Asterisks indicate statistically significant differences compared to 1-day-old females: *p ≤ 0.05, **p ≤ 0.01, ***p ≤ 0.001, ****p ≤ 0.0001.

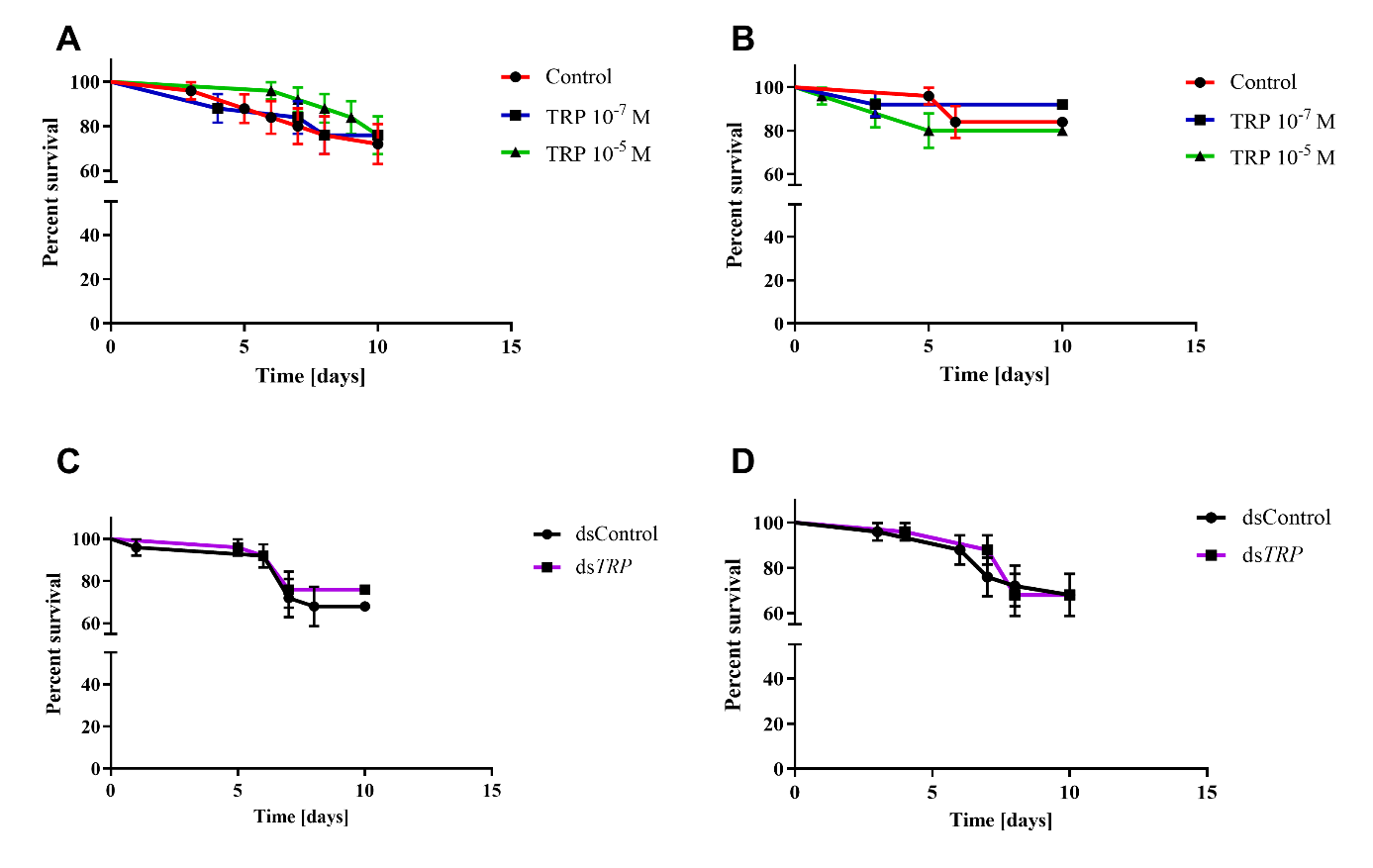

**Fig. 2S.** Survival curves of 1-day-old (A) and 4-day-old (B) *T. molitor* females after injection of physiological saline (control, red line) or Tenmo-TRP-7 at a concentration 10^-7^ M (blue line) and 10^-5^ M (green line). Additionally survival was also tested after ds*GmLys* (dsControl, black line) and ds*TRP* (violet line) application in 1-day-old (C) and 4-day-old (D) females. The values are presented as the means ± SEs. Statistical comparison of estimated survival curves based on the Gehan-Breslow-Wilcoxon test; ns – nonsignificant difference.

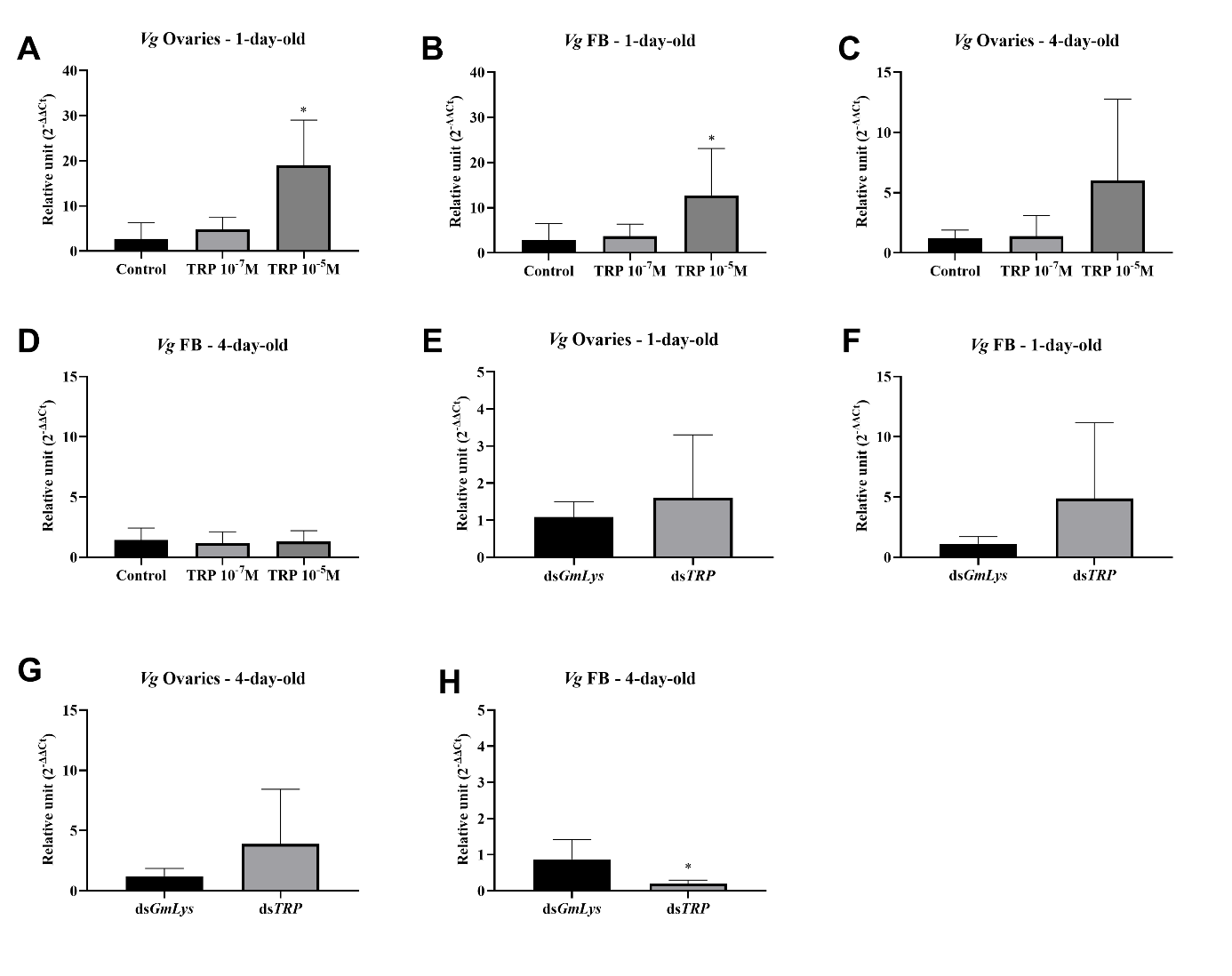

**Fig. 3S**. Changes in the expression levels of genes encoding vitellogenin (Vg) after application of Tenmo-TRP-7 at a concentration of 10^-7^ M and 10^-5^ M (A-D) and dsRNA directed against *Galeria melonella* lysozyme (ds*GmLys*) and TRP precursor (ds*TRP*) (E-H). The changes were examined in 1-day-old females in the ovaries (A and E), fat body (B and F), and 4-day-old females in the ovaries (C and G) and fat body (D and H). Values are presented as means ± SD. Asterisks indicate statistically significant differences compared to control or ds*GmLys* (for RNAi experiment), *p ≤ 0.05.

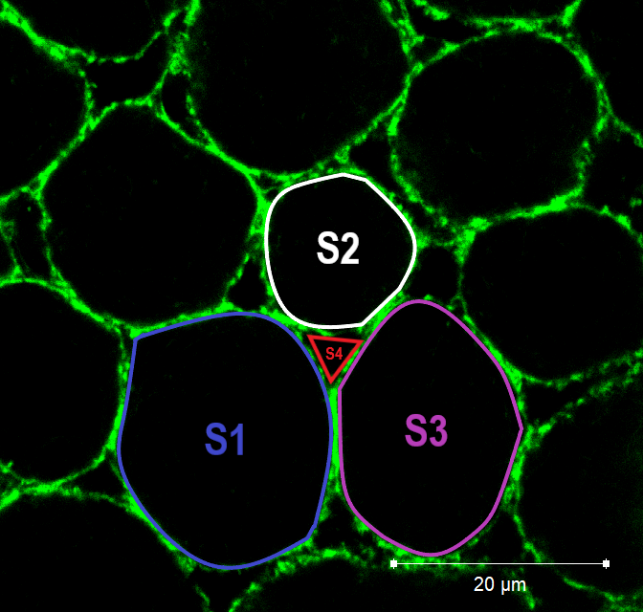

**Fig. 4S.** Illustration of the follicular epithelium and the approach to its analysis. The patency index: P = S₄ / (S₁ + S₂ + S₃), where S₄ is the surface area of the intercellular space, calculated as the average area of all visible intercellular spaces in a given experimental variant; S₁, S₂, and S₃ are the surface areas of follicular cells adjacent to a given intercellular space, calculated as the average area of these cells from the microscope image.

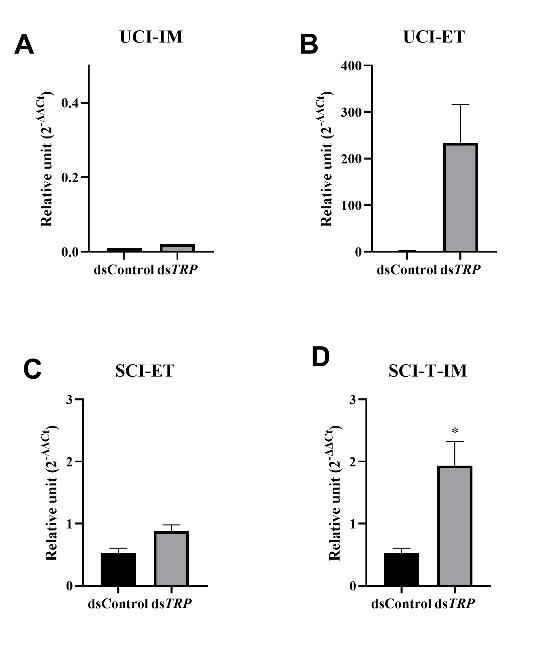

**Fig. 5S.** Efficiency of TRP knockdown on the 8^th^ day after topical application. dsControl – The mRNA quantity of TRP was measured relative to *Galeria mellonella* lysozyme (GmLys) as a negative control using RT-qPCR. Changes were analyzed in females with unsclerotized cuticle, to whom dsRNA was applied to the intersegmental membrane (A, UCI-IM) and the epidermis under the elytra (B, UCI-ET), as well as in females with sclerotized cuticle, to whom dsRNA was applied to the epidermis under the elytra (C, SCI-ET) and a mixture of dsRNA and 0.1% Triton-X to the intersegmental membrane (D, SCI-T-IM). The values are presented as the means ± SDs, *p ≤ 0.05.
